## Supplementary Material for "Upc2-mediated mechanisms of azole resistance in *Candida auris*"

**Table S1: Plasmids used in this study**

| Name | Description | Reference |
| --- | --- | --- |
| pJK795 | *NatR* selection marker for nourseothricin | (26) |
| pYM70 | *HygR* selection marker for hygromycin | (27) |
| Clp-p*ACT1*-3xFLAG-MNase-SV40-*CYC-SAT1* | *CauNi* and SAT1 selection marker for hygromycin | (28) |
| pjli6 | $P_{ADH1}$, *CauNi* and *SAT1* selection marker for nourseothricin | This study |
| pjli7 | $P_{ADH1}$, *MRR1-3XHA* Tag, *CauNi* and *SAT1* selection marker for nourseothricin | This study |
| pjli8 | $P_{ADH1}$, *MRR1-3XHA* Tag, $T_{ACT1}$, *CauNi* and *SAT1* selection marker for nourseothricin | This study |
| pjli11 | $P_{ADH1}$, *TAC1b-3XHA* Tag, $T_{ACT1}$, *CauNi* and *SAT1* selection marker for nourseothricin | This study |
| pjli12 | $P_{ADH1}$, *UPC2-3XHA* Tag, $T_{ACT1}$, *CauNi* and *SAT1* selection marker for nourseothricin | This study |

**Table S2: Primers and RNAs guides used in this study**

| Primer | Aim | Sequence (5' → 3') |
|---|---|---|
| ADH1p_PF_KpnI | *ADH1* promotor amplification containing KpnI restriction site to clone pjli6 | ACACTGGTACCCGAGATAGATCGAAATACGCTC |
| ADH1p_PR_KasI_NheI | *ADH1* promotor amplification containing NheI restriction site to clone pjli6 | ACACTGCTAGCGGCGCCGATTTCGTGAAGATTGATTGATGATG |
| MRR1_PF_KasI | *MRR1* amplification containing KasI restriction site to clone pjli7 | ACACTGGCGCCATGGTATCTTCGAAAGATC TGGC |
| MRR1_PR_BsrGI_Tag_CS_NruI_NheI | *MRR1* amplification containing BsrGI restriction site, the 3xHA Tag sequence, codon stop of *MRR1*, and NruI, NheI restriction sites to clone pjli7 | ACACTGCTAGCTCGCGATTAAGCGTAATCCGGAACATCGTATGGGTAAGCGTAATCCGGAACATCGTATGGGTAAGCGTAATCCGGAACATCGTATGGGTATGTACACACATCAAGCATCTCTTCGAATG |
| ACT1t_PF_NheI | *ACT1* terminator amplification containing NheI restriction site to clone pjli8 | ACACTGCTAGCTCTCTCAAAGATGGTTAGTATTCTTGC |
| ACT1t_PR_NheI | *ACT1* terminator amplification containing NheI restriction site to clone pjli8 | ACACTGCTAGCGAGTGACTGCTGGGTAGTAG |
| TAC1b_PF_KasI | *TAC1b* amplification containing KasI restriction site to clone pjli11 | ACA CTG GCG CCA TGA GTG CAG TGG TGA AGG G |

| | | |
|---|---|---|
| TAC1b_PR_BsrGI | *TAC1b* amplification containing BsrGI restriction site to clone pjli11 | ACACTTGTACAAAGCCCATTATCGAAGAAG AAATTAG |
| UPC2_PF_KasI | *UPC2* amplification containing KasI restriction site to clone pjli12 | ACACTGGCGCCATGTCTATGAAAGAAGAGC AACAGC |
| UPC2_PR_BsrGI | *UPC2* amplification containing BsrGI restriction site to clone pjli12 | ACACTTGTACAAATATCGTTAGACCCAATGAACCC |
| pjli8_ADH1_PF | pjli8, pjli11 and pjli12 sequencing / *TAC1b*, *MRR1*, *UPC2* hyperactivation verification | AGCAACACCGGTGGAATTTCC |
| MRR1_PrimerF2 | pjli8 sequencing | GCTAAGCAGATGTTCGATGCG |
| MRR1_PrimerF3 | pjli8 sequencing | CTCATCCATTTCTGGGAAGCC |
| MRR1_PrimerF4 | pjli8 sequencing | GAATAAGCATACTTTGTCCTTGGG |
| MRR1_PrimerF5 | pjli8 sequencing | TACGTCTTTATCACACTTATGCTCC |
| MRR1_PrimerF6 | pjli8 sequencing / *MRR1* deletion verification | ATCTTTCACTCAAGACATTTTTGATCTC |
| MRR1_PrimerF7 | pjli8 sequencing | AAGTCAAGGAGTTGGTAACCATTTG |

| | | |
|---|---|---|
| TAC1_PrimerF2 | pjli11 sequencing | AGCTGAGTCTCCAGGAAACTG |
| TAC1_PrimerF3 | pjli11 sequencing | GAAGCAGTACAAGTTTTTGAACTCG |
| TAC1_PrimerF4 | pjli11 sequencing | GCTGGTAAACATCGTCGACG |
| TAC1_PrimerF5 | pjli11 sequencing | CTTCGCCATGAACTGGATAATTAC |
| TAC1_PrimerF6 | pjli11 sequencing | AAGAATGACCCTACACTTTCCAAC |
| UPC2_PF2 | pjli12 sequencing | CCGCCACCTAACCTATCAAAC |
| UPC2_PF3 | pjli12 sequencing / *UPC2* deletion verification | GCCAGGTTTCCAACACGATTAC |
| UPC2_PF4 | pjli12 sequencing | GCTGAAGTCATTTGCTGATCTCG |
| CauNi_verif_PF | *TAC1b*, *MRR1*, *UPC2* hyperactivation verification | TGGCCTCTTGATAAGGGCTG |
| sat1_verif_PR | *TAC1b*, *MRR1*, *UPC2* hyperactivation verification | GAGACTGTGCGCGACTCC |
| pjli8_ACT1t_PR | *TAC1b*, *MRR1*, *UPC2* hyperactivation verification | GTACGCTAAAGGGTCATGAGC |
| UPC2_del_PF1 | Construction of *UPC2* deletion cassette | CTATCCGTGATAGCTAAGTCGG |

| | | |
|---|---|---|
| UPC2_del_PR1 | Construction of *UPC2* deletion cassette | GTATTCTGGGCCTCCATGTCCTCGGGAGAGTTGAATCCTC |
| UPC2_del_PF2 | Construction of *UPC2* deletion cassette | GAGGATTCAACTCTCCCGAGGACATGGAGGCCCAGAATAC |
| UPC2_del_PR2 | Construction of *UPC2* deletion cassette | CCTTCACTACTAACTCTTCACACTCAGTATAGCGACCAGCATTCAC |
| UPC2_del_PF3 | Construction of *UPC2* deletion cassette | GTGAATGCTGGTCGCTATACTGAGTGTGAA GAGTTAGTAGTGAAGG |
| UPC2_del_PR3 | Construction of *UPC2* deletion cassette | AGAGGGGTTGCAAAGGAGAG |
| UPC2_del_PF4 | Construction of *UPC2* deletion cassette | CATGCAGGTGTAGGTTACGAAAG |
| UPC2_del_PR4 | Construction of *UPC2* deletion cassette | TCGTGGTGGAGATTCTCATGG |
| UPC2_del_PR1(hygro) | Construction of *UPC2* deletion cassette | TATCGAACAGCAAGCACTATACCTCGGGAG AGTTGAATCCTC |
| UPC2_del_PF2(hygro) | Construction of *UPC2* deletion cassette | GAGGATTCAACTCTCCCGAGGTATAGTGCT TGCTGTTCGATA |
| UPC2_del_PR2(hygro) | Construction of *UPC2* deletion cassette | CTTCACTACTAACTCTTCACACTATTTTATGATGGAATGAATGGGATG |
| UPC2_del_PF3(hygro) | Construction of *UPC2* deletion cassette | CATCCCATTCATTCCATCATAAAATAGTGTGAAGAGTTAGTAGTGAAG |
| NAT1_743_F | *UPC2* deletion verification | AGTTCTTCGTTTCACTGAGTATACG |
| UPC2_del_verif_PR | *UPC2* deletion verification | CGAGGTAGTGGAAGGAGAAG |

| | | |
|---|---|---|
| F444Lverif_PF | *UPC2*, *MRR1*, *MDR1* deletion verification | CCACCCAAGGCATTTCTATATC |
| MRR1del_PF1 | *MRR1* deletion | CGCTCCACTCTTAGAAAAATGGTC |
| MRR1del_PR1 | *MRR1* deletion | TATCGAACAGCAAGCACTATACGAGAAGTGTTAATTGCCGCT |
| MRR1del_PF2 | *MRR1* deletion | AGCGGCAATTAACACTTCTCGTATAGTGCTTGCTGTTCGATA |
| MRR1del_PR2 | *MRR1* deletion | CGCATGTATATTTACGTAGTATGATTTTATGATGGAATGAATGGGATG |
| MRR1del_PF3 | *MRR1* deletion | CATCCCATTCATTCCATCATAAAATCATACTACGTAAATATACATGCG |
| MRR1del_PR3 | *MRR1* deletion | TTGGTCATAAACTCAATATCCCTCC |
| MRR1del_PF4 | *MRR1* deletion | GCATCTTCCATGACTAGACGC |
| MRR1del_PR4 | *MRR1* deletion | CATAGTCAGGTAGAGAGCCTTC |
| MRR1del_Verif_PR2 | *MRR1* deletion verification | CACCAGATTGGAGTTCGGTAAG |
| MRR1_PrimerR2 | *MRR1* deletion verification | TTGTGGGTGTTATCGGTGCC |
| MDR1del_PF1 | *MDR1* deletion | AAATAGGCAGGCGGGCAACG |
| MDR1del_PR1 | *MDR1* deletion | TATCGAACAGCAAGCACTATACGTGGAGATTGAAGATGCGTTG |
| MDR1del_PF2 | *MDR1* deletion | CAACGCATCTTCAATCTCCACGTATAGTGCTTGCTGTTCGATA |
| MDR1del_PR2 | *MDR1* deletion | GATCAGGGCTACATCGTCTTATTTTATGATGGAATGAATGGGATG |
| MDR1del_PF3 | *MDR1* deletion | CATCCCATTCATTCCATCATAAAATAAGACGATGTAGCCCTGATC |
| MDR1del_PR3 | *MDR1* deletion | GGAGGTTTCATCCGCAAAAGC |
| MDR1del_PF4 | *MDR1* deletion | CCGGTGGTGGTTCACGTAAG |
| MDR1del_PR4 | *MDR1* deletion | CGGCTGCGAAAAGAGGTTCC |
| MDR1del_verif_PR | *MDR1* deletion verification | CGCTGCCAAAGTGCTAATATCC |
| MDR1_verif_PF | *MDR1* deletion verification | GTCTGTCAAGGCTGACTACC |
| ACT1_F | *ACT1* RT_PCR | GAAGGAGATCACTGCTTTAGCC |
| ACT1_R | *ACT1* RT_PCR | GAGCCACCAATCCACACAG |
| UPC2_PF(sybrgreen) | *UPC2* RT_PCR | AGGGTGTGTCACAGGTGTTG |

| | | |
|---|---|---|
| UPC2_PR(sybrgreen) | *UPC2* RT_PCR | CGCCTCCCAAGAAATCAAGC |
| ERG11_PF(sybrgreen) | *ERG11* RT_PCR | GTTTGCCTACGTGCAATTGG |
| ERG11_PR(sybrgreen) | *ERG11* RT_PCR | GTAGTCGACTGGTGGAAGCG |
| TAC1b_PF(sybrgreen) | *TAC1b* RT_PCR | AGAAGTGAACGCTCCTTCCG |
| TAC1b_PR(sybrgreen) | *TAC1b* RT_PCR | GGCGTGAATTTCCTGTCGTC |
| CDR1_F | *CDR1* RT_PCR | GAAATCTTGCACTTCCAGCCC |
| CDR1_R | *CDR1* RT_PCR | CATCAAGCAAGTAGCCACCG |
| MRR1_PF(sybrgreen) | *MRR1* RT_PCR | TAGAGCCACTACCATCCGAC |
| MRR1_PR(sybrgreen) | *MRR1* RT_PCR | AGTTTGTGCCCTGCCTGATG |
| MDR1_F | *MDR1* RT_PCR | GAAGTATGATGGCGGGTG |
| MDR1_R | *MDR1* deletion verification/*MDR1* RT_PCR | CCCAAGAGAGACGAGCCC |
| CauNi_sg5’ | Guide RNA for *TAC1b*, *MRR1*, *UPC2* hyperactivation | CCCGGAGAUACACGGCGCCG |
| CauNi_sg3’ | Guide RNA for *TAC1b*, *MRR1*, *UPC2* hyperactivation | GCUGCAAAAUAAGGCCAGAG |
| UPC2_del_sg5’ | Guide RNA for *UPC2* deletion | AAUCUUCUGAGAAAAAGAGA |
| UPC2_del_sg3’ | Guide RNA for *UPC2* deletion | AGCAACUUGGACAUCAUGCA |
| MRR1del_Guide_RNA_5' | Guide RNA for *MRR1* deletion | UUGACGCGAGAGAAUUUGGA |
| MRR1del_Guide_RNA_3' | Guide RNA for *MRR1* deletion | ACGCCUGGAGAAUCACUAGA |
| MDR1del_sg5' | Guide RNA for *MDR1* deletion | CAAAGUGUUCACAUACCCCG |
| MDR1del_sg3' | Guide RNA for *MDR1* deletion | CCAGUAUUAUUCUACCUCAA |

**Supplementary Figure 1**

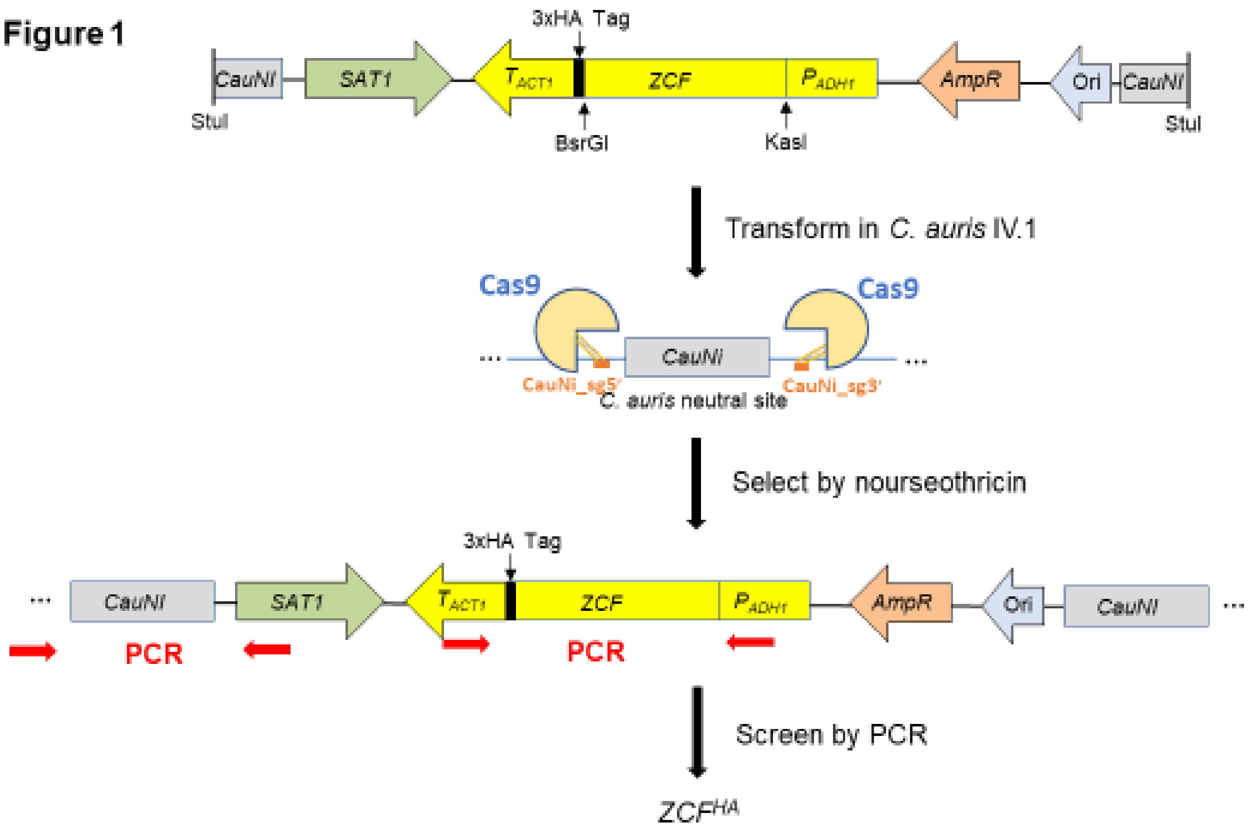


**Schematic view of the zinc cluster transcription factor (ZCF) hyperactivation system in *C. auris*.** The plasmid containing the promotor $P_{ADH1}$, the nucleotide sequence of ZCF fused by 3xHa Tag, the terminator of $T_{ACT1}$, the SAT1 cassette (nourseothricin resistance) and the *C. auris* neutral site *CauNI* was linearized by StuI. The restriction sites KasI and BsrGI were used to insert the ZCF nucleotide sequences. The linearized plasmid was transformed via electroporation in the wild-type IV.1 strain with CRISPR-Cas9 method, by targeting the upstream and the downstream regions of *CauNI* with the nucleotide-specific guide RNAs CauNi_sg5' and CauNi_sg3'.

### Supplementary Figure 2A

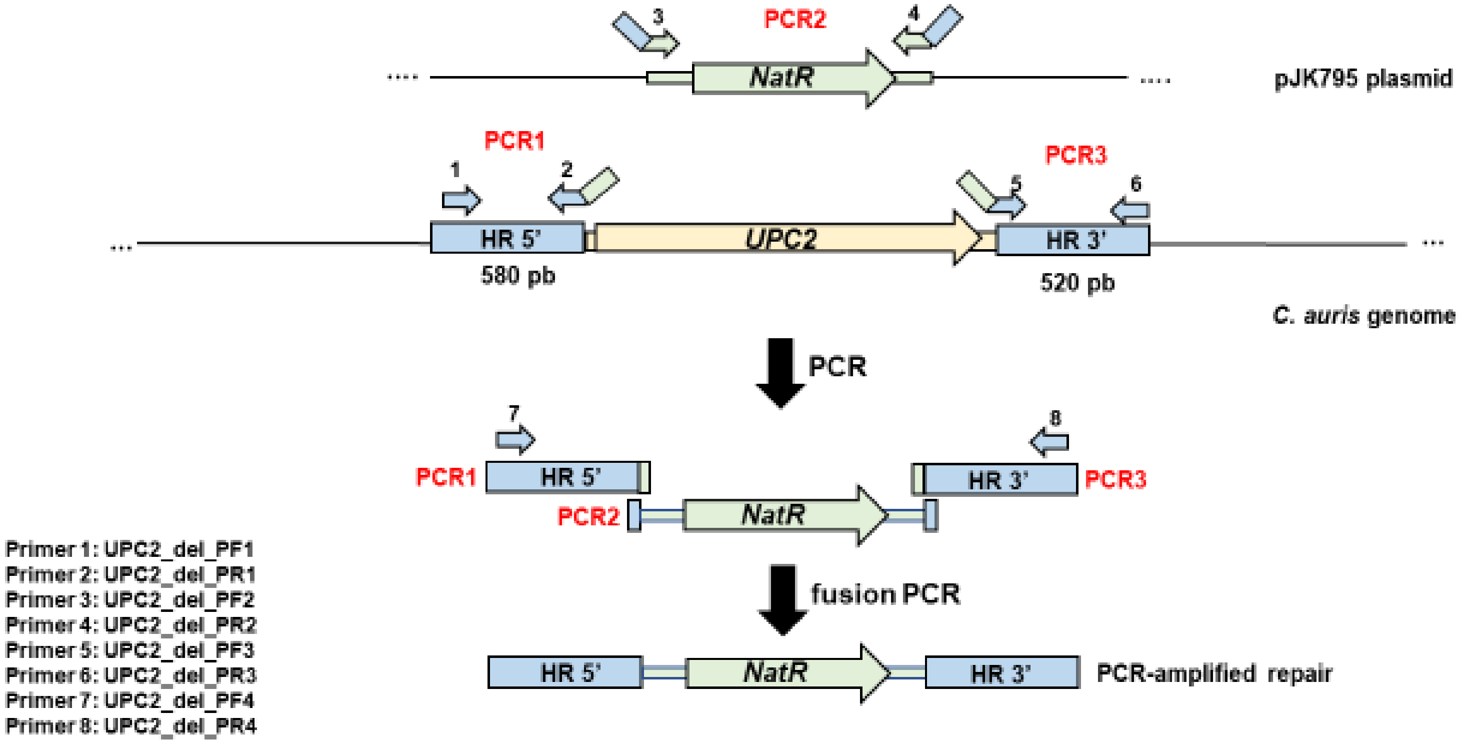


**Schematic view of the fusion PCRs for *UPC2* deletion in the IV.1 strain to generate the *upc2*Δ strain.** The fragment contains the *NatR* cassette (nourseothricin resistance) and 500 pb flanking sequences (HR 5' and HR 3') of *UPC2*. The fusion PCRs were carried out with overlapping primers as indicated. Primer 2 is reverse complemented to primer 3, and primer 4 is reverse complemented to primer 5.

### Supplementary Figure 2B

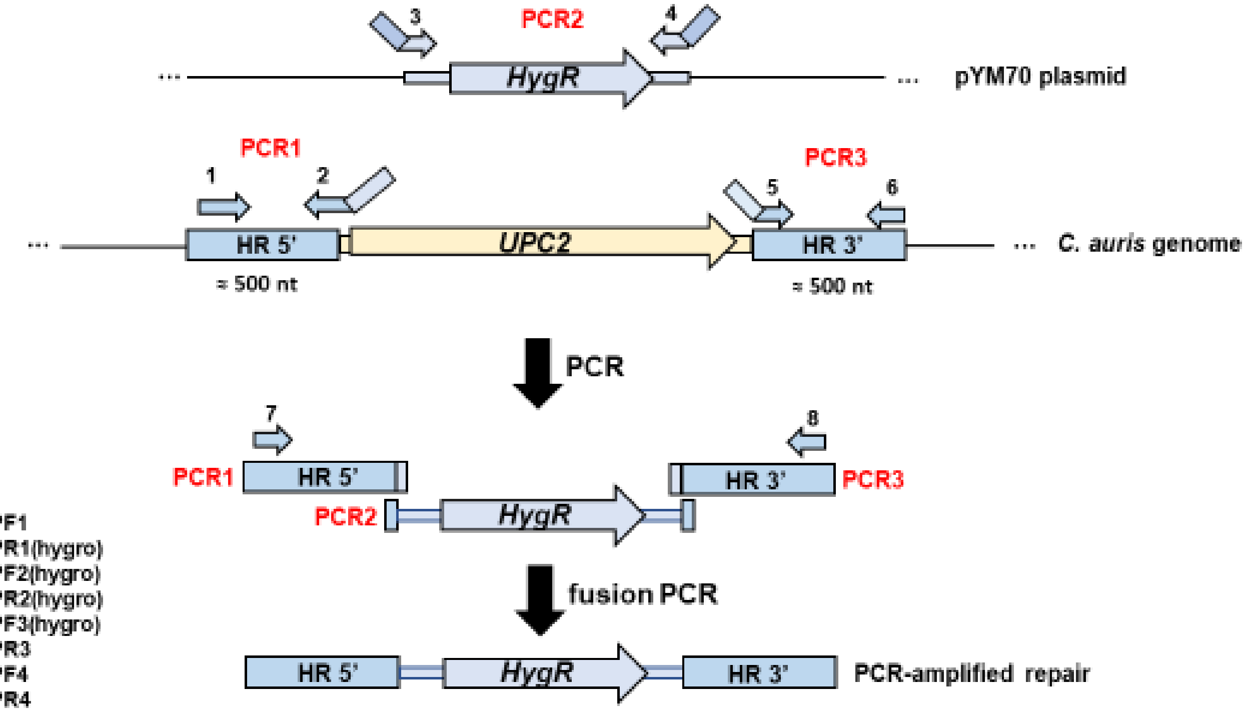


Primer 1: UPC2_del_PF1
Primer 2: UPC2_del_PR1(hygro)
Primer 3: UPC2_del_PF2(hygro)
Primer 4: UPC2_del_PR2(hygro)
Primer 5: UPC2_del_PF3(hygro)
Primer 6: UPC2_del_PR3
Primer 7: UPC2_del_PF4
Primer 8: UPC2_del_PR4

**Schematic view of the fusion PCRs for *UPC2* deletion in the *TAC1b*$^{HA}$ and *MRR1*$^{HA}$ strains to generate the *TAC1b*$^{HA}$/*upc2*Δ and *MRR1*$^{HA}$/*upc2*Δ strains.** The fragment contains the *HygR* cassette (hygromycin resistance) and 500 pb flanking sequences (HR 5' and HR 3') of *UPC2*. The fusion PCRs were carried out with overlapping primers as indicated. Primer 2 is reverse complemented to primer 3, and primer 4 is reverse complemented to primer 5.

### Supplementary Figure 3

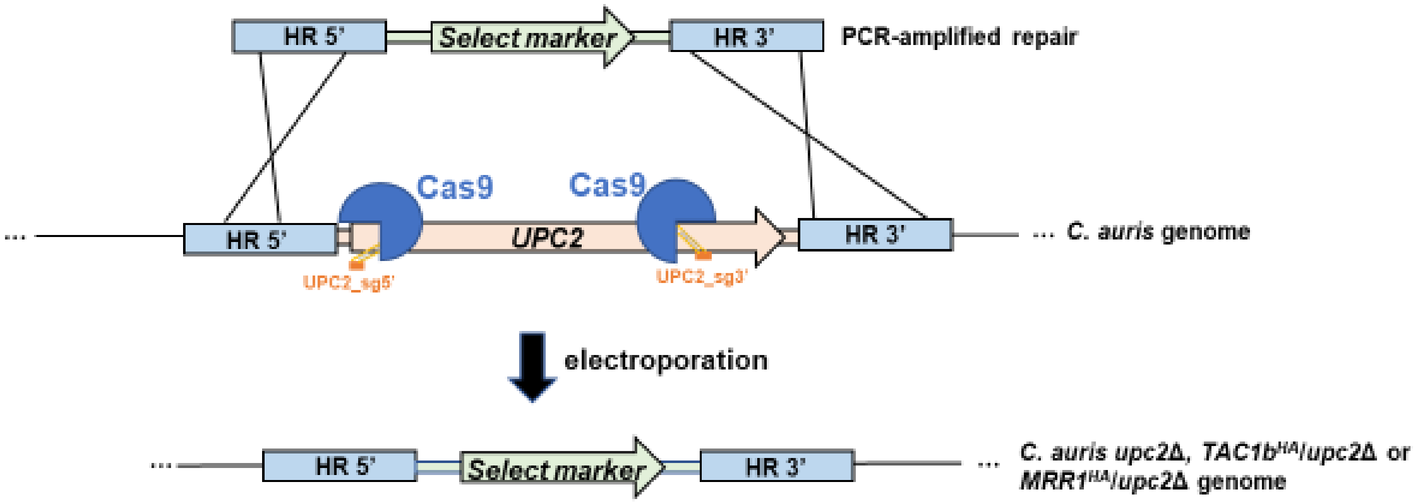


**Principle for CRISPR-Cas9-mediated deletion of *UPC2* in the IV.1, *TAC1b*$^{HA}$ and *MRR1*$^{HA}$ strains.** Two guide-RNAs were used: UPC2_sg5' and UPC2_sg3'. The selection markers *NatR* (nourseothricin resistance) and *HygR* (hygromycin resistance) were used for *UPC2* deletion in strain IV.1 and in strains *TAC1b*$^{HA}$ and *MRR1*$^{HA}$, respectively.

### Supplementary Figure 4A

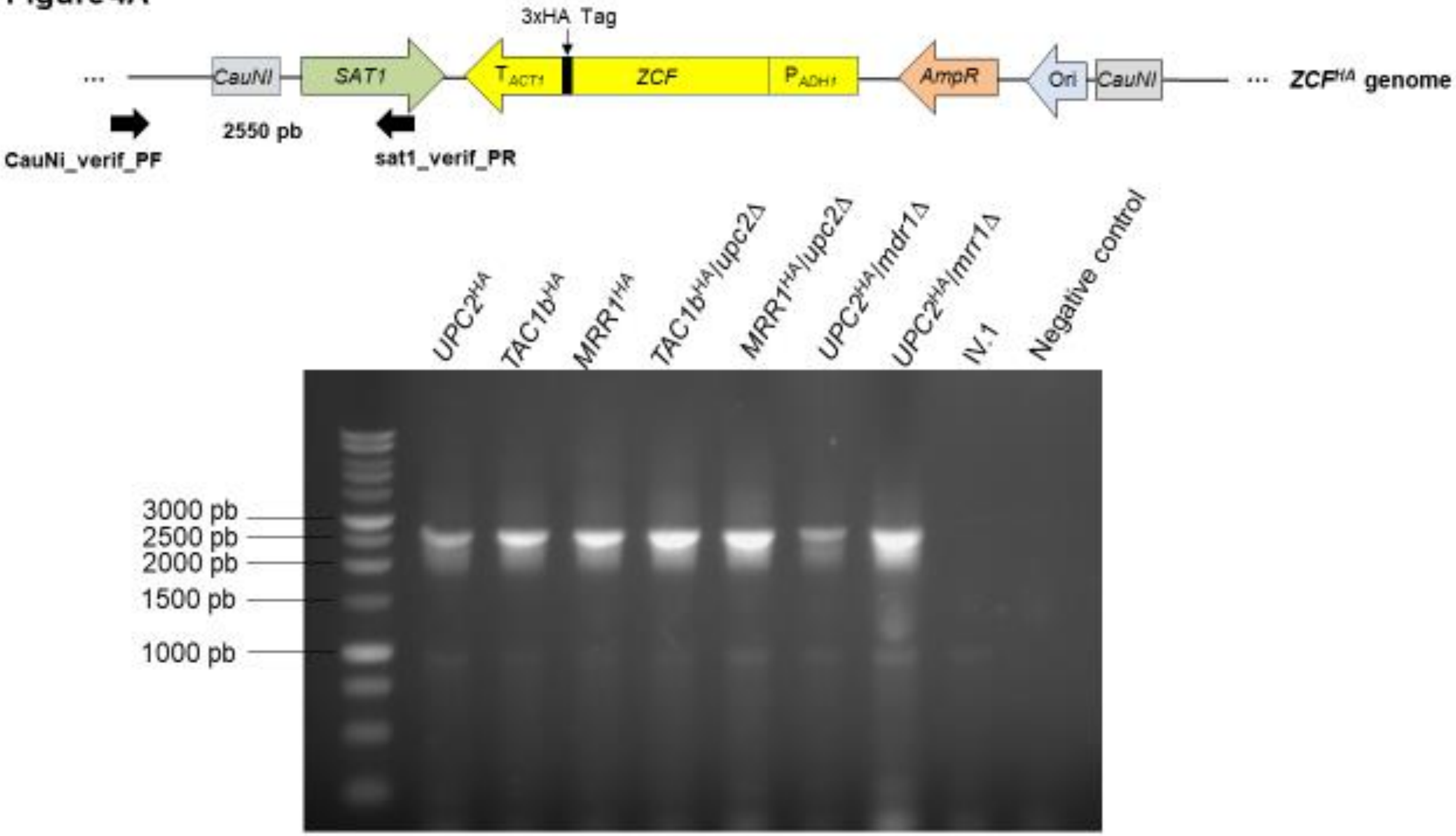


**Verification of correct integration of the transformation cassette for zinc cluster transcription factor hyperactivation in the *UPC2*$^{HA}$, *TAC1b*$^{HA}$, *MRR1*$^{HA}$, *TAC1b*$^{HA}$*/upc2*Δ, *MRR1*$^{HA}$*/upc2*Δ, *UPC2*$^{HA}$*/mdr1*Δ and *UPC2*$^{HA}$*/mrr1*Δ strains.** An approximately 2.5 kb fragment was amplified with one primer in *SAT1* and another primer in the upstream region of the *C. auris* neutral site *CauNI*. As shown on the electrophoresis gel, the PCR product was present in the mutant strains and absent in the IV.1 strain.

**Supplementary Figure 4B**

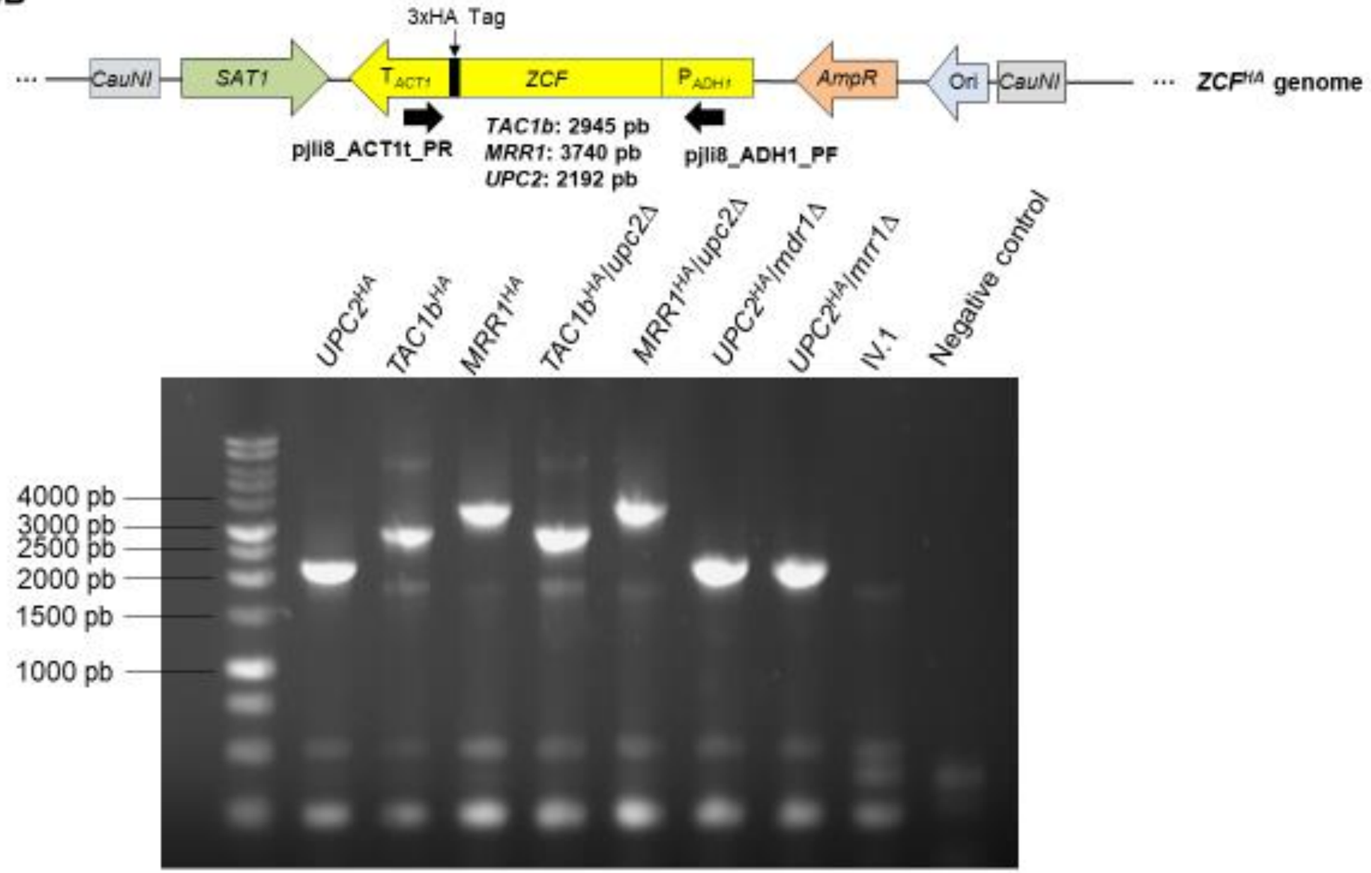


**Verification of correct integration of the transformation cassette for zinc cluster transcription factor hyperactivation in the *UPC2*$^{HA}$, *TAC1b*$^{HA}$, *MRR1*$^{HA}$, *TAC1b*$^{HA}$*/upc2*Δ, *MRR1*$^{HA}$*/upc2*Δ, *UPC2*$^{HA}$*/mdr1*Δ and *UPC2*$^{HA}$*/mrr1*Δ strains.** A fragment was amplified with one primer in the promoter $P_{ADH1}$ and another primer in the terminator $T_{ACT1}$. As shown on the electrophoresis gel, PCR products of different sizes (depending on the length of the target genes) were obtained in all the mutant strains.

### Supplementary Figure 5A

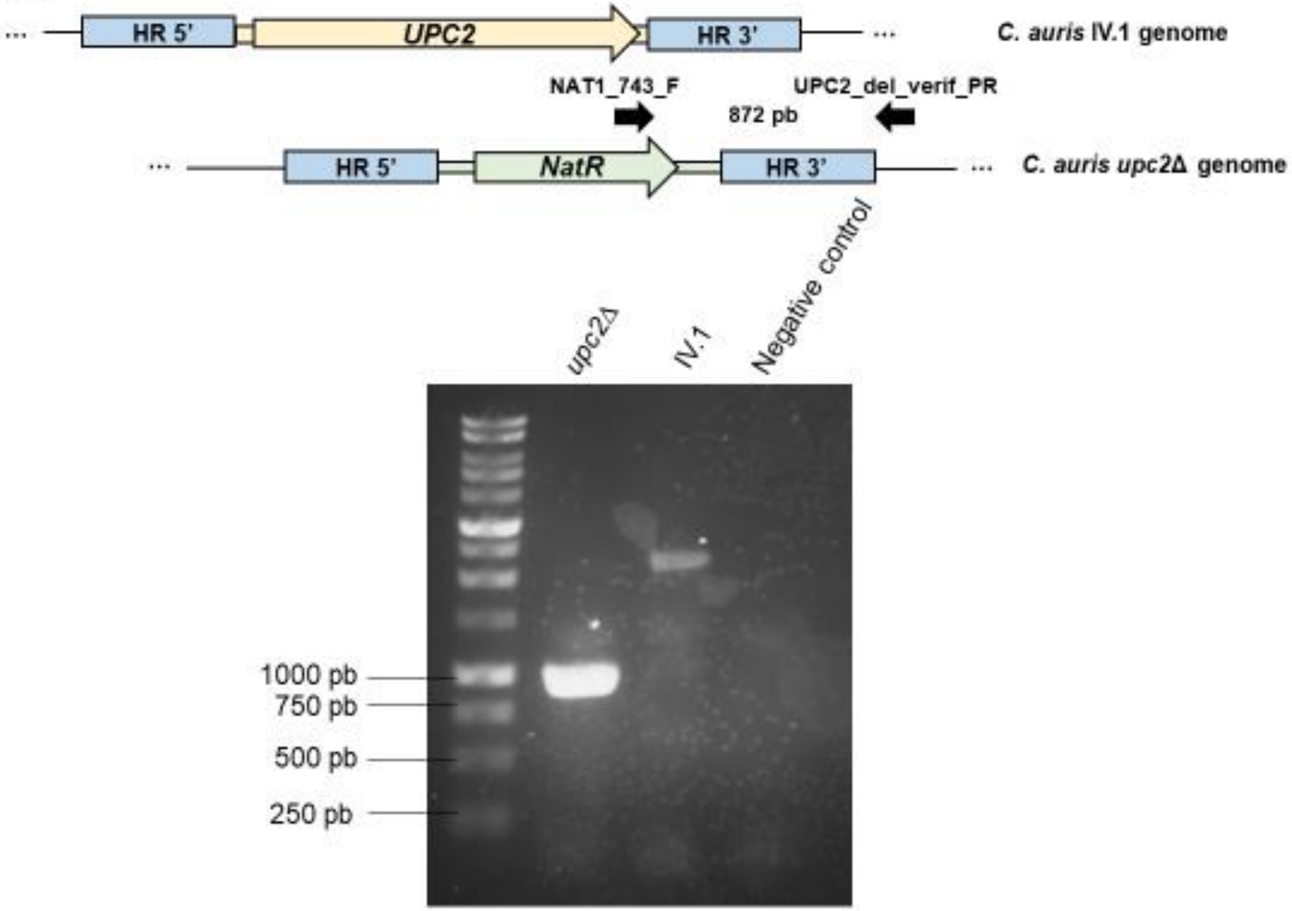


**Verification of the deletion of *UPC2* in the *upc2*Δ strain.** An approximately 900 pb fragment was amplified with one primer in *NatR* and another primer in the downstream region of HR 3'. As shown on the electrophoresis gel, the PCR product was present in the mutant strain and absent in the IV.1 strain.

Supplementary Figure 5B

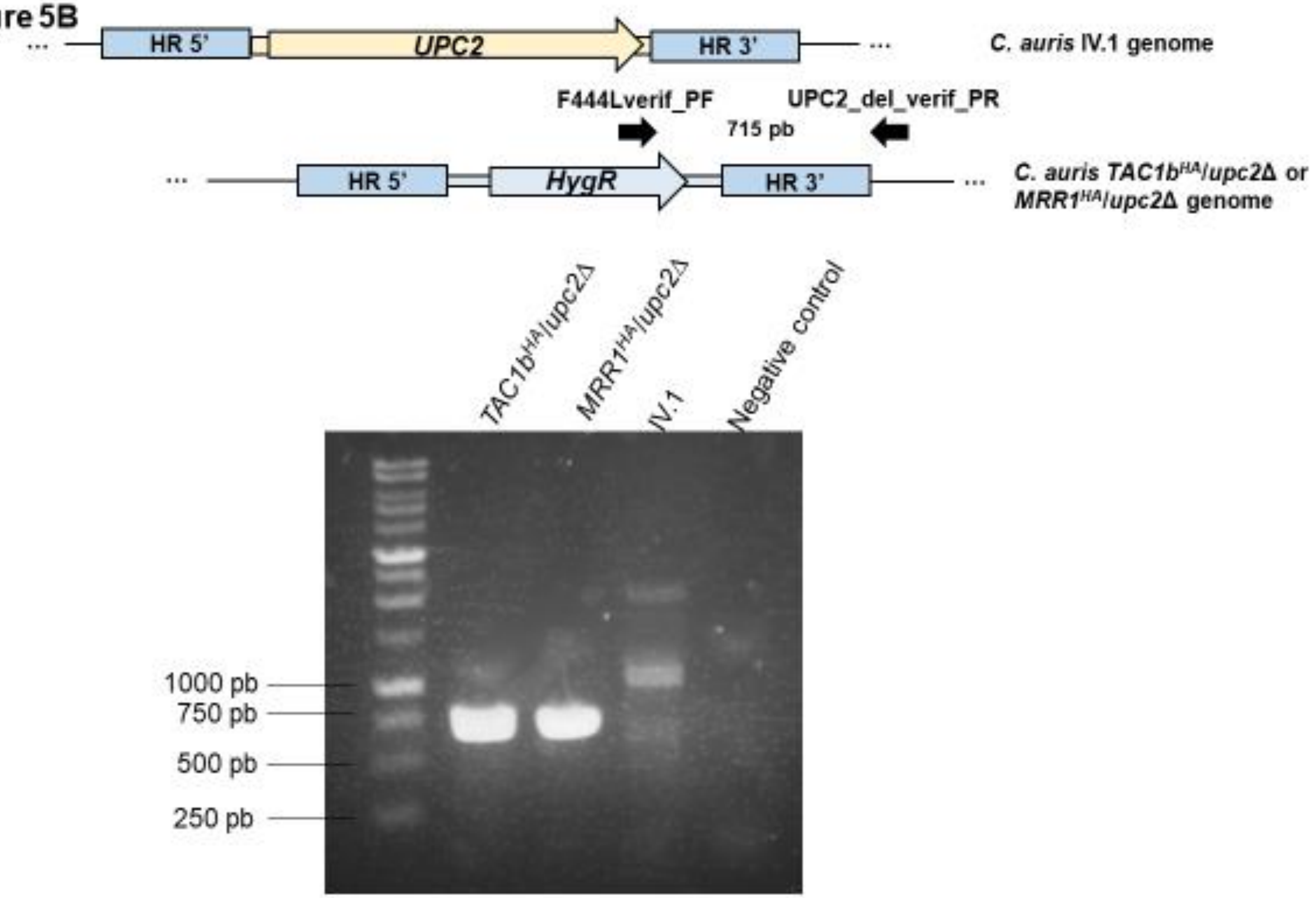


**Verification of the deletion of *UPC2* in the*TAC1b$^{HA}$/upc2Δ* and *MRR1$^{HA}$/upc2Δ* strains.** An approximately 700 pb fragment was amplified with one primer in *HygR* and another primer in the downstream region of HR 3'. As shown on the electrophoresis gel, the PCR product was present in the mutant strains and absent in the IV.1 strain.

**Supplementary Figure 5C**

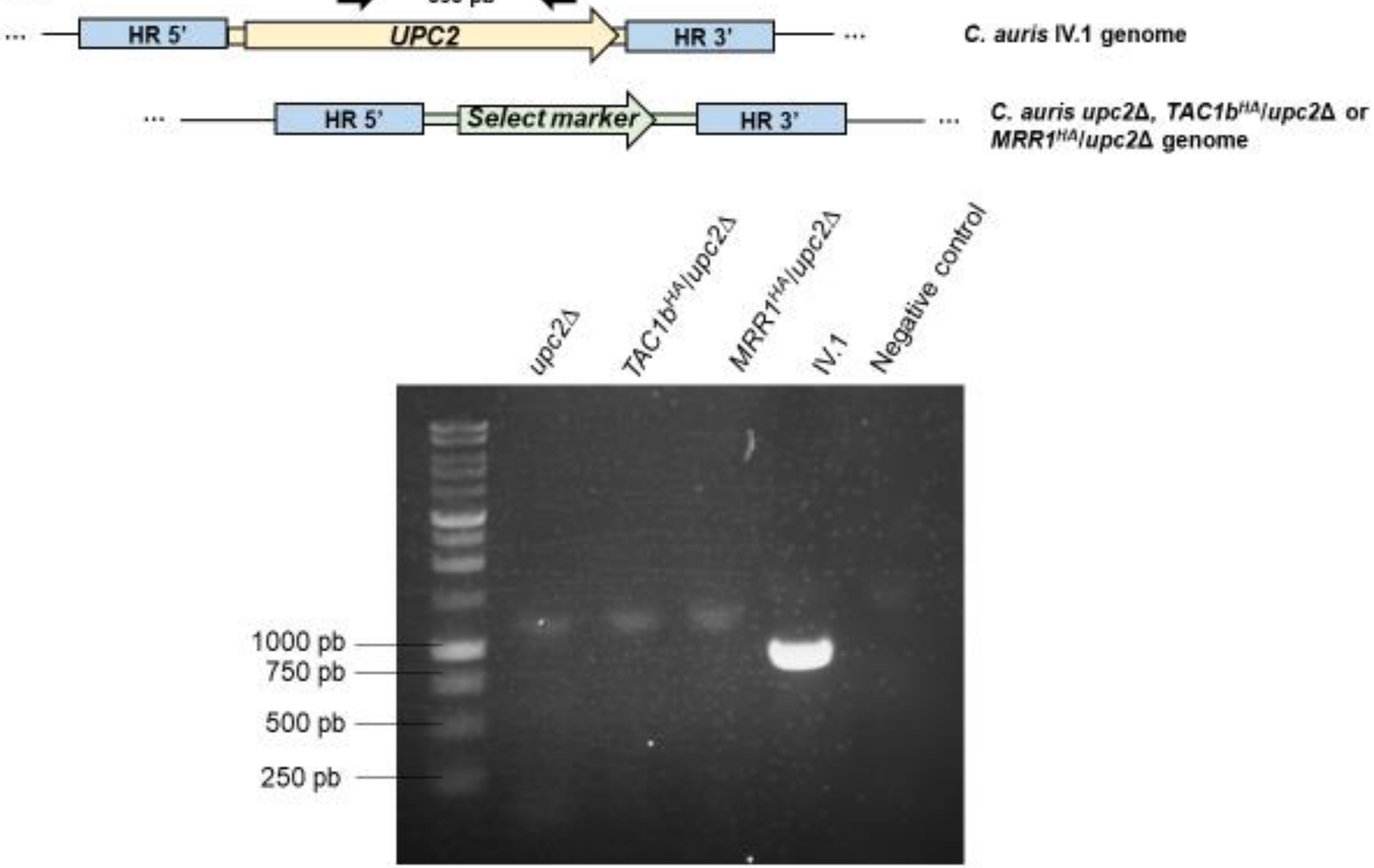


**Verification of the deletion of *UPC2* in the *upc2*Δ, *TAC1b*$^{HA}$/*upc2*Δ and *MRR1*$^{HA}$/*upc2*Δ strains.** An approximately 800 pb fragment was amplified with two primers located within *UPC2*. As shown on the electrophoresis gel, the PCR product was present in the IV.1 strain and absent in the mutant strains.

### Supplementary Figure 6A

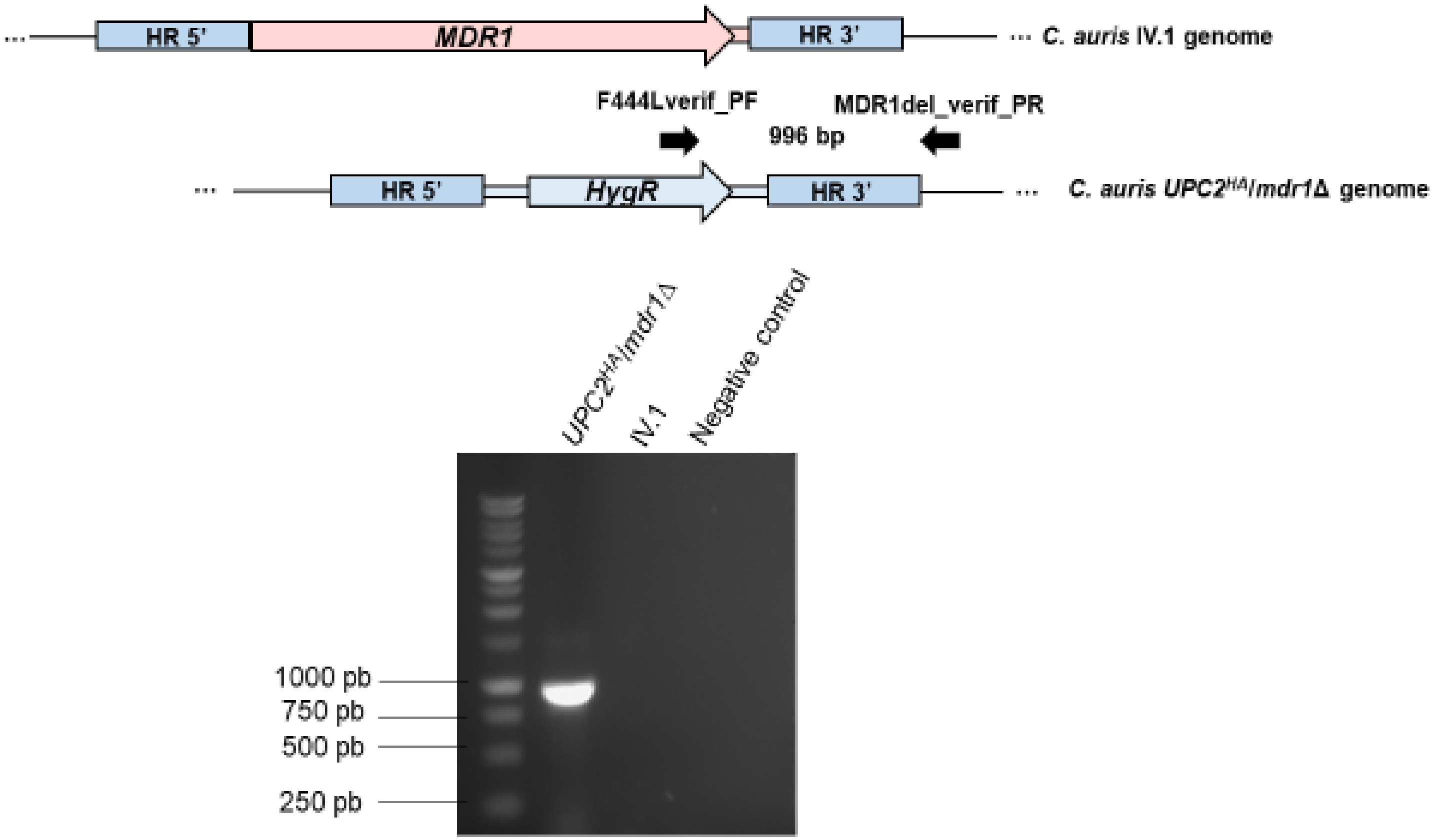


**Verification of the deletion of *MDR1* in the *UPC2$^{HA}$/mdr1Δ* strain.** An approximately 1 kb fragment was amplified with one primer in *HygR* and another primer in the downstream region of HR 3'. As shown on the electrophoresis gel, the PCR product was present in the mutant strain and absent in the IV.1 strain.

**Supplementary Figure 6B**

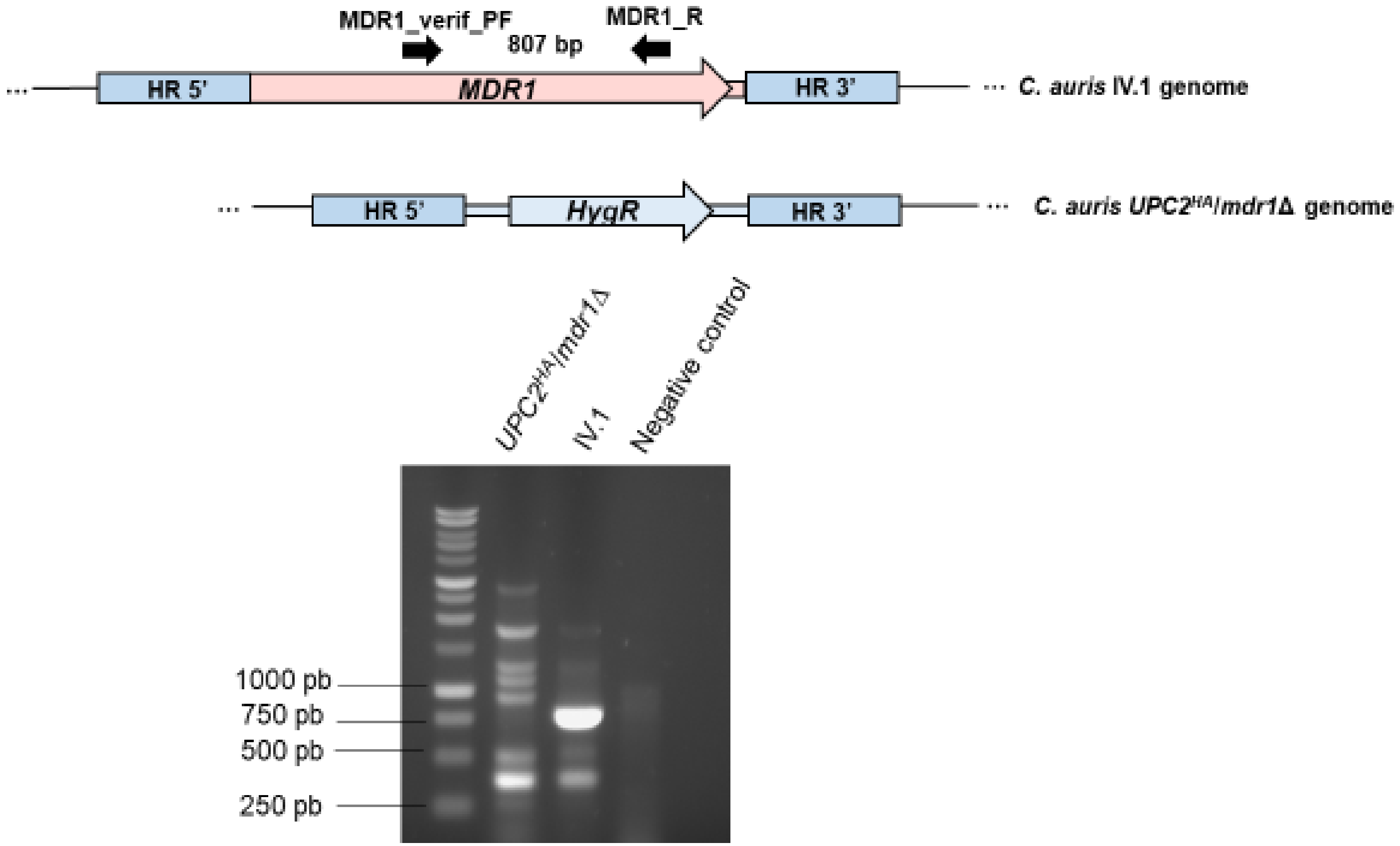


**Verification of the deletion of *MDR1* in the *UPC2$^{HA}$/mdr1Δ* strain.** An approximately 800 pb fragment was amplified with two primers located within *MDR1*. As shown on the electrophoresis gel, the PCR product was present in the IV.1 strain and absent in the mutant strain.

### Supplementary Figure 7A

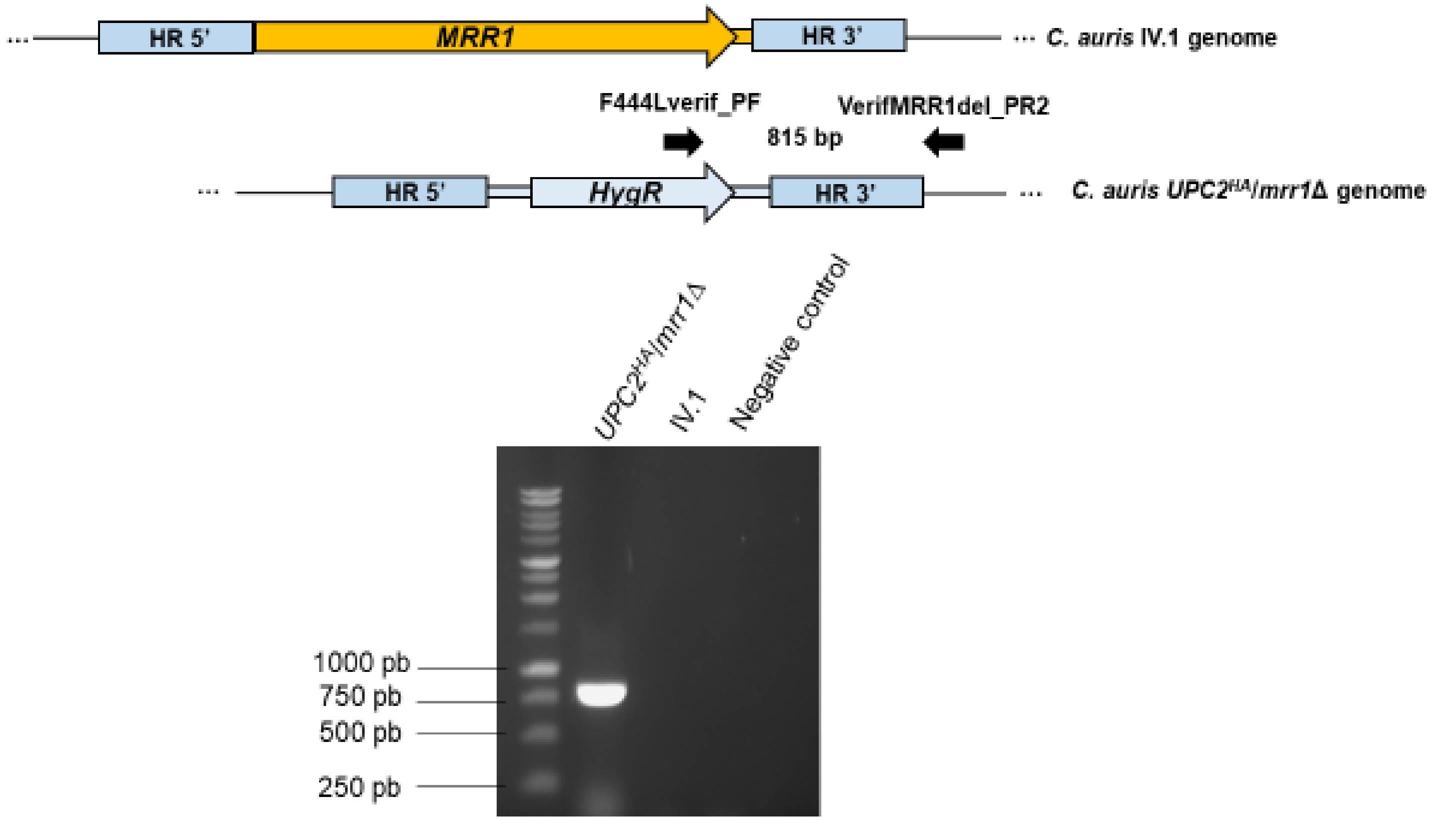


**Verification of the deletion of *MRR1* in the *UPC2$^{HA}$/mrr1Δ* strain.** An approximately 800 pb fragment was amplified with one primer in *HygR* and another primer in the downstream region of HR 3'. As shown on the electrophoresis gel, the PCR product was present in the mutant strain and absent in the IV.1 strain.

### Supplementary Figure 7B

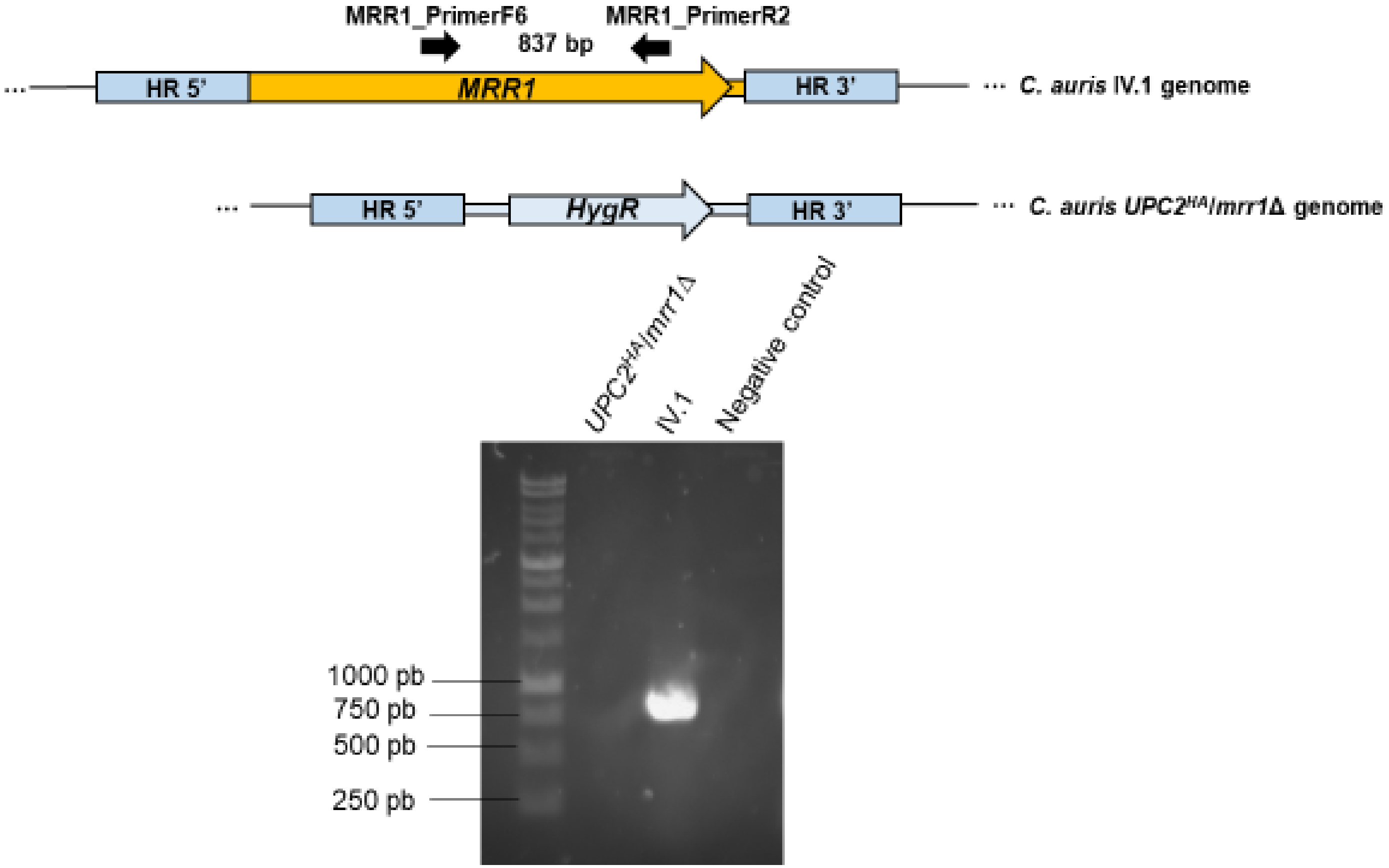


**Verification of the deletion of *MRR1* in the *UPC2$^{HA}$/mrr1Δ* strain.** An approximately 800 pb fragment was amplified with two primers located within *MRR1*. As shown on the electrophoresis gel, the PCR product was present in the IV.1 strain and absent in the mutant strain.
